# Functional heterogeneity in the fermentation capabilities of the healthy human gut microbiota

**DOI:** 10.1101/2020.01.17.910638

**Authors:** Thomas Gurry, Le Thanh Tu Nguyen, Xiaoqian Yu, Eric J Alm

## Abstract

The human gut microbiota is known for its highly heterogeneous composition across different individuals. However, relatively little is known about functional differences in its ability to ferment complex polysaccharides. Through *ex vivo* measurements from healthy human donors, we show that individuals vary markedly in their microbial metabolic phenotypes (MMPs), mirroring differences in their microbiota composition, and resulting in the production of different quantities and proportions of Short Chain Fatty Acids (SCFAs) from the same inputs. We also show that aspects of these MMPs can be predicted from composition using 16S rRNA sequencing. From experiments performed using the same dietary fibers *in vivo*, we demonstrate that an ingested bolus of fiber is almost entirely consumed by the microbiota upon passage. We leverage our *ex vivo* data to construct a model of SCFA production and absorption *in vivo*, and argue that inter-individual differences in quantities of absorbed SCFA are directly related to differences in production. Taken together, these data suggest that personalized dietary fiber supplementation based on an individual’s MMP is an attractive therapeutic strategy for treating diseases associated with SCFA production.

## Introduction

The symbiotic relationship between host and gut microbiota is intimately related to host diet. For example, ruminants derive the majority of their caloric intake from the microbial fermentation of indigestible polysaccharides in their diet. In humans, only a fraction of total caloric intake is derived from fermentation of dietary fibers and other Microbiota Accessible Carbohydrates (MACs), but the resulting metabolites also play other important physiological roles (Davie, 2003; Iraporda et al., 2015; Kuo, 2013). In particular, the Short Chain Fatty Acids (SCFAs) acetate, propionate and butyrate, which are the major by-products of the microbial fermentation of dietary fibers in the gut, exert a number of forces on the host’s physiology. While it has long been known that butyrate serves as the dominant energy source for colonocytes (Roediger, 1980, 1982), and is therefore critically important to maintaining a healthy gut, it has been shown more recently that imbalances in SCFA production can be associated with disease. In the case of microbial acetate, increased production and turnover have been shown to activate glucose-stimulated insulin secretion via a gut-brain axis mediated process, which can result in insulin resistance and subsequent obesity, in addition to increasing a patient’s risk of developing Type 2 diabetes (Perry et al., 2016). In contrast, propionate and butyrate activate gluconeogenesis, with beneficial effects to host metabolism and the regulation of plasma glucose levels (De Vadder et al., 2014; Pingitore et al., 2017; Zhao et al., 2018). Indeed, increased butyrate production has been causally linked to improved insulin response after an oral glucose-tolerance test, whereas abnormalities in propionate production were associated with an increased risk of Type-2 Diabetes (Sanna et al., 2019).

Many of these effects are mediated by short chain fatty acid receptors in the gut epithelium. However, a critical property of colonic SCFAs is their activity as histone deacetylase (HDAC) inhibitors (Davie, 2003). This ability to regulate gene expression in host cells has associated SCFAs with a growing list of clinical indications. Butyrate, in particular, has been proposed to exert anti-inflammatory pressure on the host’s immune system through several mechanisms, which include differentiation of regulatory and IL-10-producing T cells, down regulation of IL-6 production, pro-inflammatory T cell apoptosis, and suppression of IFN-γ-mediated inflammation in the colonic epithelium (Zimmerman et al., 2012). Consistent with these findings, when Clostridia strains were computationally ranked by regulatory T cell induction capability, the highest ranked strains were predicted to produce significantly higher butyrate than the lower ranked strains (Stein et al., 2018). These data complement measured associations between Inflammatory Bowel Disease (IBD) and depletions in butyrate-producing organisms (e.g. *Faecalibacterium prausnitzii*) (Machiels et al., 2013) to indicate that SCFAs likely play important roles in the disease etiology. Moreover, the fact that SCFAs produced by the microbiota in the colon can be absorbed into the blood stream suggests that their effects on gene expression may transcend the gut and affect distal tissues in ways that currently poorly understood. It is therefore of significant clinical interest to improve our quantitative understanding of SCFA production in the gut, in order that it may be modulated towards a particular clinical outcome.

Several different biochemical fermentation pathways in the gut metagenome result in SCFAs as a final product. These pathways begin with the hydrolysis of complex dietary polysaccharides to their constituent oligo- and monosaccharides by members of the microbiota encoding the appropriate Polysaccharide Utilization Loci (PULs) and Glycoside Hydrolases (GHs), which differ widely between different bacterial species (Sonnenburg et al., 2010) and even strains within a species (Filippis et al., 2019). These monosaccharides can then be fermented by a number of pathways to ultimately result in acetate, propionate or butyrate. Importantly, there is cross-talk between different pathways: for example, the most prevalent butyrate-producing pathway in the human gut involves the enzyme butyryl-CoA : acetate-CoA transferase (Duncan et al., 2002), which exchanges a butyrate moiety for an acetate moiety on the CoA molecule and releases free butyrate. A similar enzyme exists for propionate in one of its three main production pathways (Reichardt et al., 2014). Thus, the pool of available acetate affects the production of both propionate and butyrate. Other products of bacterial fermentation can also serve as intermediates in SCFA production: lactate, in particular, can act as a substrate for further fermentation into acetate, propionate and butyrate. Moreover, the SCFA-producing organisms may not be able to directly ferment a specific polysaccharide themselves; instead, they rely on the preliminary degradation of these dietary fibers into hexoses and pentoses by other bacteria encoding these carbohydrate active enzymes. This gives rise to a host of cooperative microbial networks which act in concert to produce the overall SCFA profile present in an individual’s colon. For example, lactic acid-utilizing, butyrate-producing bacteria (e.g. *Eubacterium hallii* and *Anaerostipes caccae*) depend on the presence of lactic acid-producing bacteria such as *Bifidobacterium adolescentis* to produce butyrate (Belenguer et al., 2006). Thus, the combination of dietary inputs (which specific dietary fibers, in which quantities) and the composition of an individual’s microbiota together dictate what ratios and absolute quantities of SCFAs are produced in their gut, with potentially important effects on physiology. In this study, we sought to gain a better understanding of the variation in fermentation capabilities in the microbiota of different individuals within the healthy human population. We present an *ex vivo* framework for measuring the production of SCFAs of an individual’s stool microbiota in response to challenge with specific dietary fibers. Performing these experiments *ex vivo* allows us to quantify SCFA accumulation accurately and circumvents the technical difficulties associated with measurements of SCFA production *in vivo* (invasiveness) or SCFA measurements from raw stool upon passage (unknown extent of SCFA absorption by the gut epithelium).

## Results

### Measuring a microbial metabolic phenotype *ex vivo*

We first sought to obtain *ex vivo* measurements of SCFA production in response to different dietary polysaccharides, in order to quantify the degree of heterogeneity in the fermentation capabilities of the healthy population’s gut microbiota. To do this, stool from 40 healthy human participants was homogenized into a slurry in anaerobic conditions and spiked with inulin, pectin or cellulose (cf. Methods). The slurry was then allowed to evolve over time and samples obtained at regular intervals in order to quantify SCFA concentration at each time point (Fig. 1a). In order to determine the appropriate sampling frequency, we performed pilot experiments in which we analyzed the trajectory of each SCFA concentration over a 24h period. We found that only a subset of participants appeared to converge to a final SCFA concentration prior to the 24h timepoint, but that all participants exhibited a linear production rate in the 0-4h time window (Fig. 1b). These data were in good agreement with concentrations of inulin measured from the stool over time using an inulin-specific ELISA assay: after 4h, a significant fraction of inulin substrate remained, but this was almost entirely consumed by 24h in five of the six participants tested (Fig. S3). Since we sought to perform the experiments in an environment that best mimicked conditions in the colon, we analyzed 16S rRNA data and determined that community structure or diversity were not significantly altered between timepoints 0 and 4h, with minor changes being observed between 4h and 24h (Fig. S4 and S5a). Participants’ slurry did not resemble each other more at 4h than they did at 0h, but did increase in similarity by the 24h timepoint (Fig. S6). Samples noticeably clustered by participant rather than by timepoint or condition (Fig. S5b). Moreover, changes in pH due to an accumulation of acidic SCFAs was limited to a drop from approximately neutral to 5.5 during the first four hours (Fig. S7). Together, these data informed our decision to use the linear production regime observed between 0-4h as the most appropriate period in which to measure SCFA production *ex vivo* as a proxy for SCFA production *in vivo*. As a result, we chose to measure two biological replicates for the timepoints 0, 2h and 4h, and compute production rates between timepoints 2h and 4h. There was generally good agreement between biological replicates, with an overall coefficient of variation of 23.5% and no discernible bias in any specific SCFA or at any specific timepoint (Fig. S1). Linear regressions between the sample’s Bristol stool scale and total SCFA concentration produced, as well as between the C_T_ values from the qPCR amplification of 16S rRNA from the stool, did not suggest any apparent artifact induced by stool consistency or density, arguing that a correction factor for stool consistency or microbial load was not required and samples of various consistencies could be compared to one another (Fig. S8).

**Figure 1.**
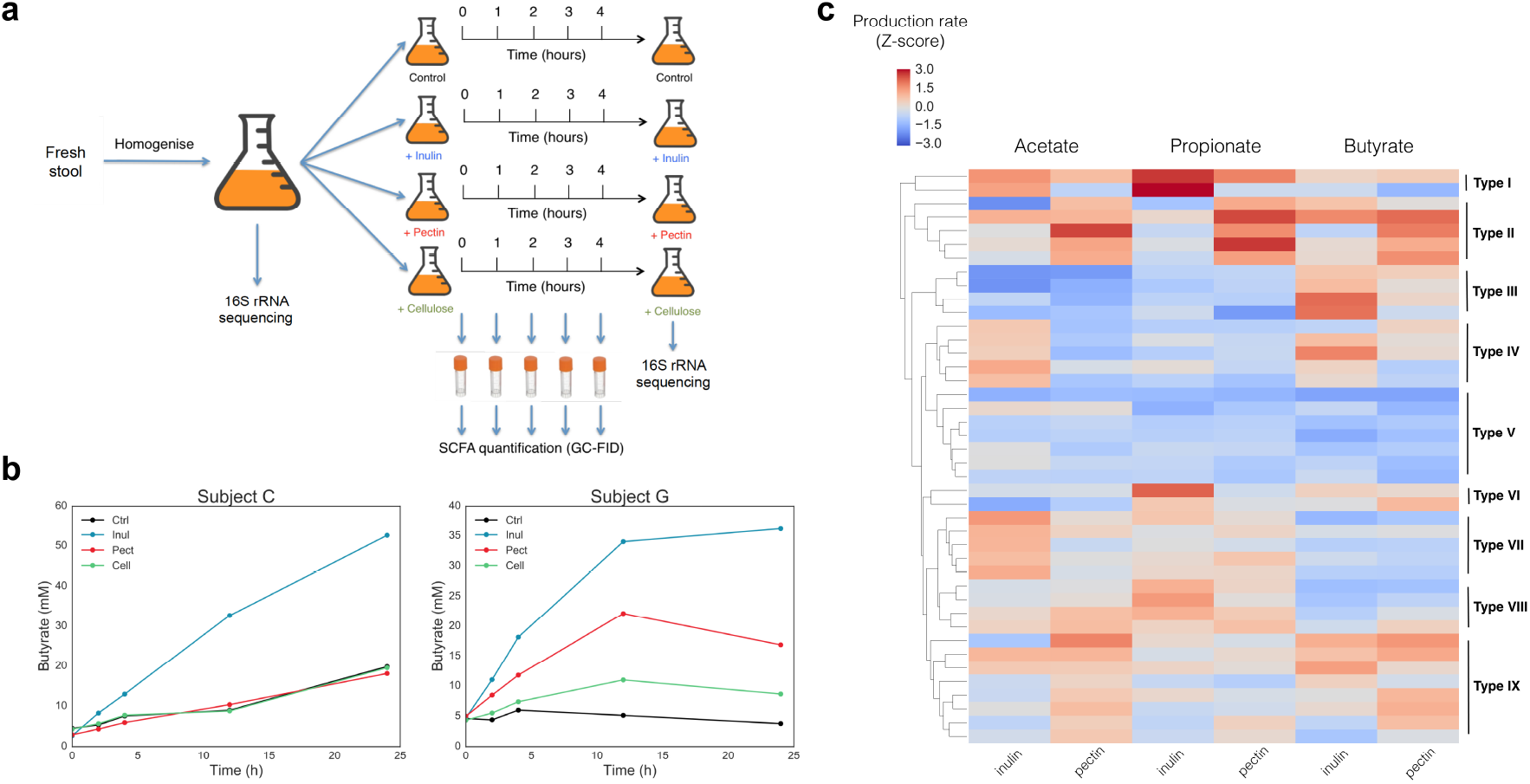
Microbial metabolic phenotype varies significantly in the healthy population. (a) Schematic of assay setup and sampling frequency. (b) 24h time traces of butyrate concentration over time in response to inulin, pectin, cellulose and control, in two different participants. Participant C only produces butyrate from inulin, while participant G produces it from pectin as well. Both participants have linear production regimes in the 0-4h window used to calculate production rate. (c) Acetate, propionate and butyrate production rates from inulin and pectin in different participants, presented as Z-scores computed across all participants. Each row represents measurements from a single sample. SCFA production rates were measured ex vivo in mM/h for each participant in response to each condition. Production rates were computed between timepoints 2h and 4h for each condition and production rates from the control condition (no spike-in) were subtracted. Cellulose timepoints were indistinguishable from control and therefore were not presented.

We found that participants differed greatly in their SCFA production profiles (Fig. 1c). From the same dietary fiber input, the different microbiotas produced significantly different quantities and ratios of SCFAs. We define the resulting SCFA production rates in response to different dietary fibers, quantified as standardized scores compared to the other participants in the dataset, as an individual’s Microbial Metabolic Phenotype (MMP). Hierarchical clustering of MMPs indicated discernible groups: for example, individuals with MMP Type I were strong producers of propionate from inulin; in contrast, participants with MMP Type II were strong producers of propionate from pectin (Fig. 1c). Thus, improving an individual’s production of a given SCFA will not necessarily rely on the same polysaccharide to reach the same effect; put differently, the same polysaccharide will have different effects in different individuals depending on their MMP. These data argue that there is significant heterogeneity in the healthy human population when it comes to functional degradation of fibers in the gut and the SCFAs produced, but that MMPs cluster into discernible types, which can be used to guide future dietary interventions.

### Predicting microbial metabolic phenotype from community composition

We then asked the question whether we could predict a participant’s MMP from community composition alone, defined here as the relative abundances of 97% *de novo* OTUs obtained from 16S rRNA sequencing of the stool prior to incubation with the different fibers. We trained Random Forest Classifiers (RFCs) to predict whether a given microbiota had a high or low production rate of a given SCFA in response to a given fiber, defined by a production rate z-score of greater than or equal to 0, or less than 0, respectively. Performance varied by SCFA, with the highest accuracies obtained in predicting butyrate production in response to inulin (AUC=0.87) and pectin (AUC=0.79) (Fig. 2a).

**Figure 2.**
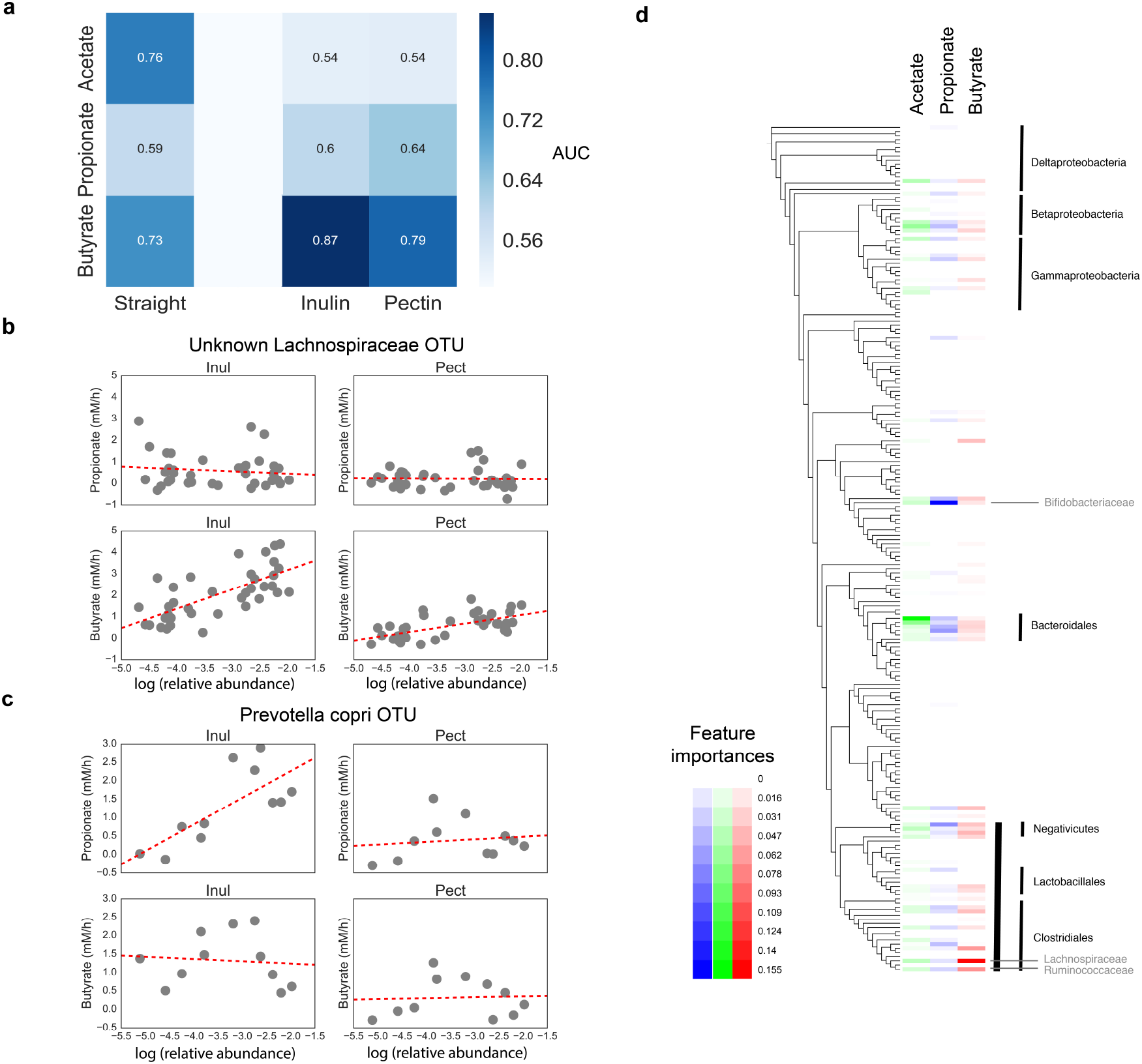
SCFA production capacity can be predicted from individual bacterial OTUs. (a) AUC values for different RFCs trained either to predict high or low SCFA content in stool at baseline, or high or low SCFA production rate *ex vivo* in response to specific dietary fibers. High and low production is defined according to the z-score across all participants in the study. (b) Relationship between propionate and butyrate production rates, and the relative abundance of an unassigned OTU of the Lachnospiraceae family, showing a relationship between its relative abundance and butyrate production in response to inulin, specifically. (c) Similar relationship but specific to the relative abundance of a *Prevotella copri* OTU and propionate production in response to inulin. (d) Feature importances from RFCs trained to predict high or low SCFA production in response to inulin from bacterial family abundances.

We also tested whether straight stool SCFA contents could be predicted from 16S rRNA sequencing and found moderate predictive power for acetate and butyrate (AUC=0.76 and AUC=0.73, respectively). Though these data indicated that community composition is somewhat predictive of the resulting stool SCFA contents, manual inspection of specific OTU features that were highly ranked in terms of importance for the inulin- and pectin-specific RFCs found these to often only be associated with SCFA production in response to that specific fiber. For example, the relative abundance of an unassigned Lachnospiraceae OTU was only associated with butyrate production in response to inulin (Fig. 2b). In contrast, a specific *Prevotella copri* OTU only appeared to be associated with propionate production from inulin (Fig. 2c). These data are consistent with the fact that members of the *Prevotella* genus are known propionate producers (Chen et al., 2017a) and vary in their polysaccharide degradation capabilities (Filippis et al., 2019). Thus, a Type I MMP (Fig. 1c) is associated with a high relative abundance of the latter *P. copri* OTU. Similarly, known butyrate-producing families (e.g. Lachnospiraceae and Ruminococcaceae) are ranked highly in importance when RFCs for high/low SCFA production in response to inulin are trained on 16S rRNA data collapsed at the family level (Fig. 2d). Though more data and whole genome characterizations are required to demonstrate these relationships rigorously, these results suggest that individual OTUs are predictive of SCFA production capability from specific polysaccharides, likely due to their specific polysaccharide degradation machinery and internal fermentation pathways.

### Stability of an individual’s MMP through time

While it is known that the gut microbiota of individuals can be relatively stable for long periods of time in the absence of large perturbations (David et al., 2014), it is unclear whether an individual’s MMP will also be similarly stable through time. We therefore repeated the experiment for eight participants at timepoints separated by at least 6 months (Fig. 3a). Though some variability between timepoints was observed, the extrema of each individual’s MMP were generally preserved (Fig. 3b). A Fisher test on the contingency table resulting from pairwise comparison of each SCFA:fiber pair at the two timepoints for all individuals indicated that this stability was statistically significant (*p*=0.003; two-tailed Fisher test). These results were consistent with the fact that MMPs are associated with the relative abundance of specific members of the microbiota: an individual with a high relative abundance of the aforementioned *P. copri* OTU is likely to retain a high relative abundance of this OTU during the six month period between timepoints compared to the general population, thus retaining their ability to produce high levels of propionate from inulin through time.

**Figure 3.**
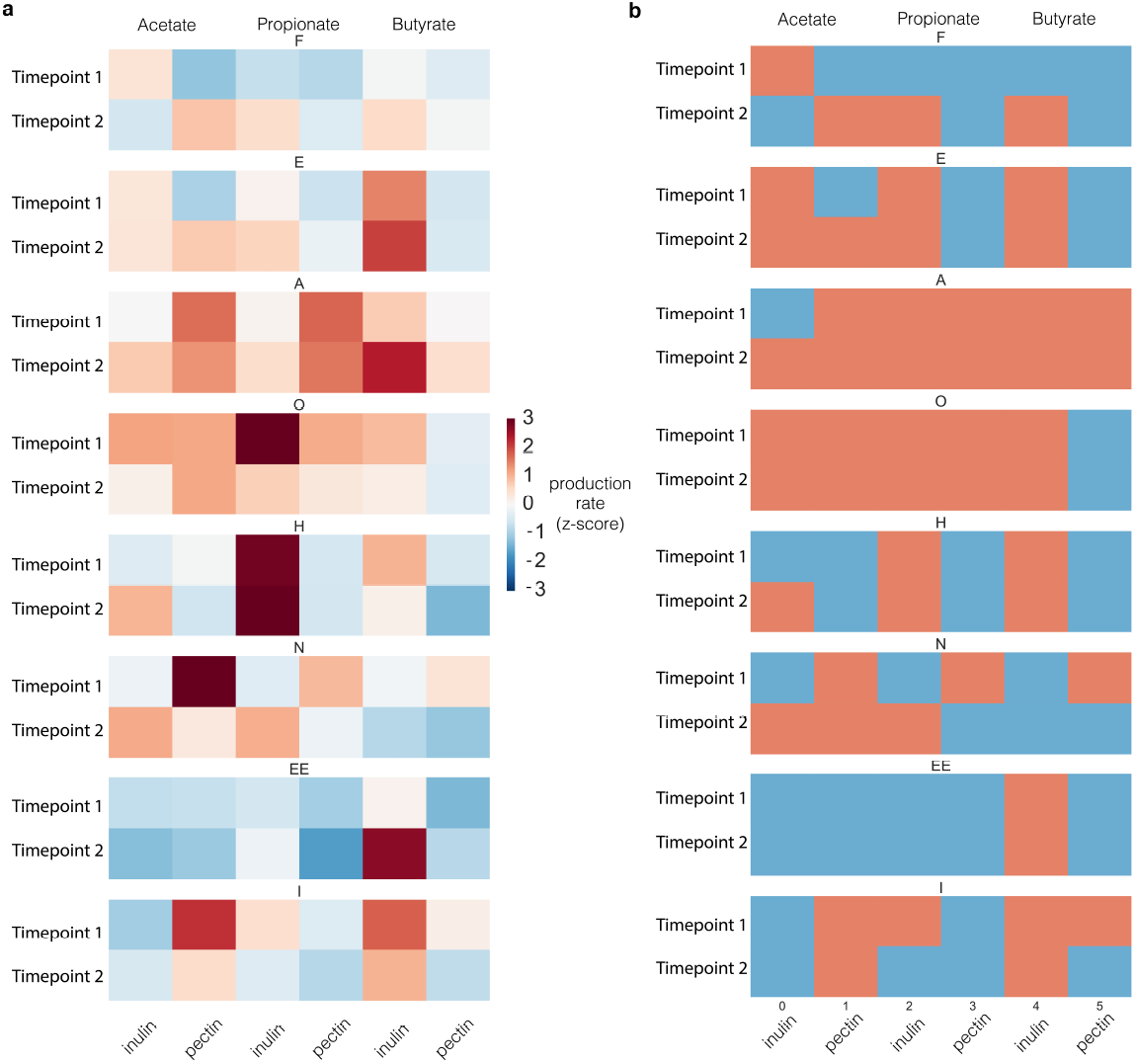
General features of an individual’s MMP are stable over time. (a) Continuous production rates of each SCFA in response to inulin and pectin for two timepoints separated by at least six months, expressed as z-scores relative to the population in the dataset. (b) The same data, but collapsed to high or low producers of a given SCFA in response to a given fiber (red = high, blue = low).

### *In vivo* model of SCFA production

In order to better understand microbial SCFA production *in vivo*, we sought to develop a quantitative model of the process (Fig. 4a). In a given participant, when a quantity of dietary fiber, *[F]*, is consumed, it is fermented into acetate, propionate and butyrate with rates that are functions of their specific microbiota. As defined, a person’s MMP is the aggregate total response of an individual’s fecal microbiota to a challenge with specific dietary fibers. The process of producing SCFAs from fiber inputs requires several steps (Fig. S9b). The first step involves breaking the fiber/polysaccharide (*F)* into smaller oligo- and monosaccharides (*O*), usually through a hydrolysis reaction encoded by extracellular enzymes (Sonnenburg et al., 2010); the second step consists of fermenting *O* into a reduced biochemical species *P* (e.g. lactate, acetate, pyruvate), which can act as a substrate for the third step, the final fermentation reaction that leads to the final product or SCFA in question. A person’s MMP is thus the bulk total of all these individual reactions (Fig. S9a).

**Figure 4.**
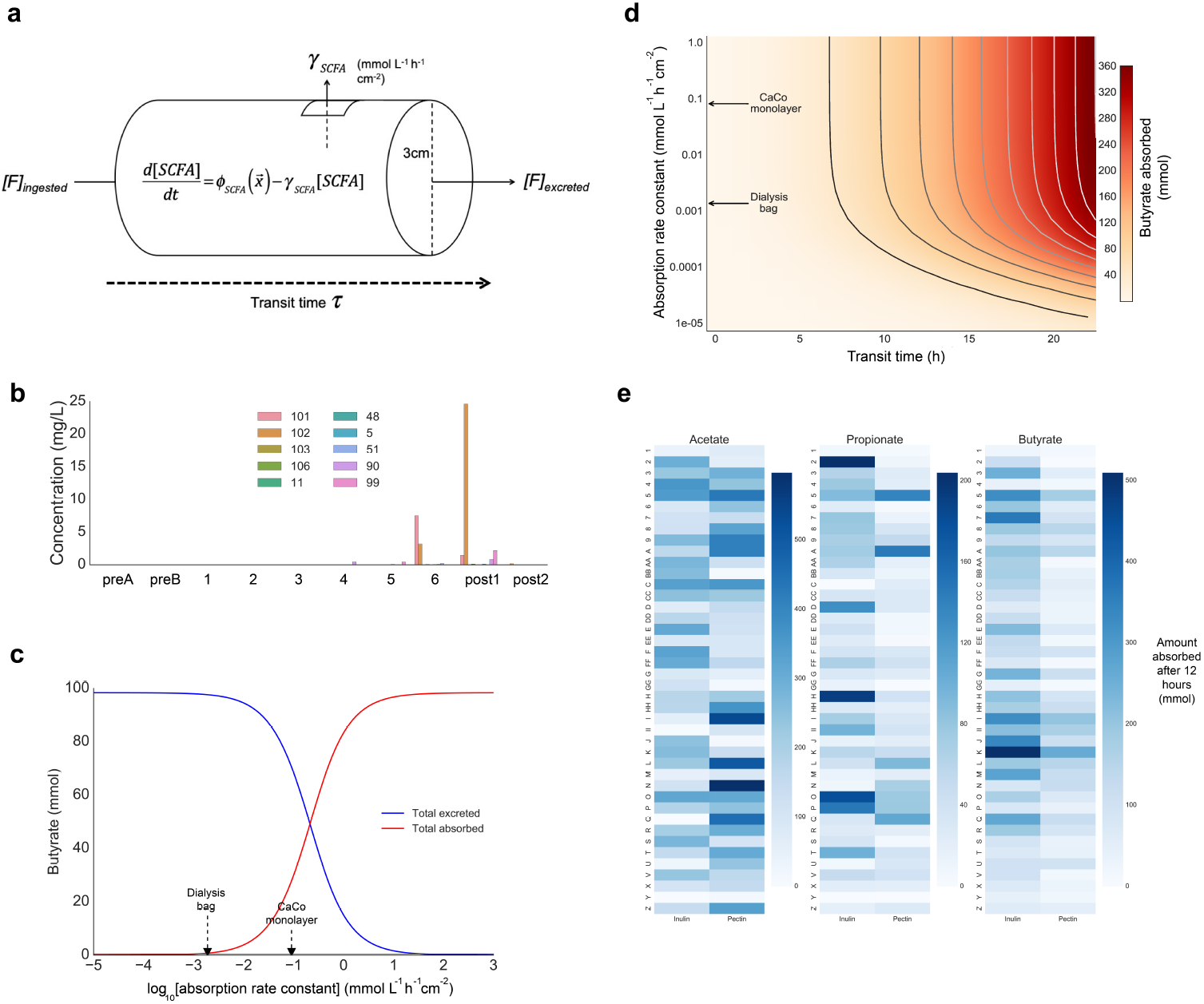
Phenomenological model of *in vivo* SCFA production predicts that inter-individual differences affects quantities absorbed. (a) Schematic of model parameters. (b) Concentration of inulin in participant stool from a previous study where participants were fed 10g of inulin on days 4, 5 and 6 against a constant (fiber-impoverished) dietary background. (c) Predicted quantities of butyrate absorbed versus excreted in the stool in participant H as a function of the colonic epithelial absorption rate constant, assuming a transit time of 12 hours. Values of the rate constant measured by the dialysis bag and CaCo monolayer approaches discussed in the text are shown explicitly. (d) Phase diagram of predicted quantity of butyrate absorbed as a function of the epithelial absorption rate constant and transit time (participant H). Values of the rate constant measured by the dialysis bag and CaCo monolayer approaches discussed in the text are shown explicitly. (e) Predicted amount of each SCFA absorbed (in mmol) using a transit time of 12 hours for each subject and the dialysis bag rate constant parameters.

In principle, some or all of the dietary fiber can be excreted in the stool without having undergone any fermentation in the colon. Thus, we sought to determine whether a consumed quantity of dietary fiber is completely fermented in the time taken for a typical bolus of food to transit through the gut. We measured fecal inulin concentrations from a previous *in vivo* study where participants were given 10g of inulin from the same vendor used in our *ex vivo* experiments as a daily supplement against a constant dietary background (Gurry et al., 2018), and found that fecal inulin concentrations ranged from undetectable to 25 mg/L on the days following inulin consumption (Fig. 4b), compared to an input concentration of approximately 10g/L (if we estimate the volume of stool in the gut to be on the order of 1L; cf. Methods). These data suggest that a bolus of inulin is almost entirely degraded by the microbiota upon passage of the stool, consistent with our measurements of inulin depletion over 24h *ex vivo* (Fig S3). We therefore assume this excretion rate of the fermentable dietary fiber is zero and that it is almost entirely consumed within the gut (i.e. *[F]*_*excreted*_ = 0).

Gut luminal concentrations of microbially-derived SCFAs is a balance between net production in the lumen and absorption by the host epithelium. Unfortunately, quantitative understanding of the rate of absorption of a given SCFA by cells in the gut epithelia is limited. Previous data collected using a dialysis bag technique suggest that SCFA absorption rate is linear with concentration at typical physiological concentrations of SCFAs (McNeil et al., 1978). Moreover, studies have shown that SCFA absorption exhibits an unexpectedly modest pH-dependence (Bugaut, 1987). We therefore modeled the absorption rate as proportional to the luminal SCFA concentration. We sought to estimate the order of magnitude of this rate constant, to better understand the fate of SCFAs produced by the microbiota in the colon (i.e. the balance between quantities absorbed versus quantities excreted). For this purpose, we used CaCo cell monolayers grown in a trans-well and measured SCFA concentrations from the media sampled from the apical and basal sides of the monolayer over a duration of 24h (cf. Methods and Supporting Information for details). Assuming the SCFA production rates measured *ex vivo* are representative of *in vivo* rates of production with the same inputs, and explicitly including the absorption rate parameters, we obtain the following system of phenomenological equations describing the time evolution of each SCFA’s concentration in the colonic lumen:

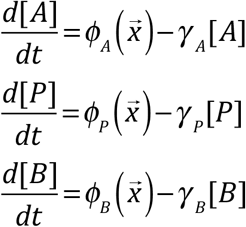

where 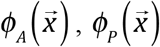 and 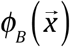 are bulk production rates of acetate, propionate and butyrate respectively, produced in response to a given input concentration of fiber *[F]*, defining an individual’s MMP.

To better understand the impact of the absorption rate constant on quantities of SCFA absorbed by the gut epithelium, we used our model to calculate the amount of butyrate absorbed in a given participant as a function of this rate constant:

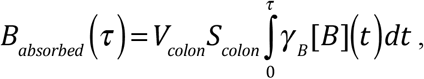

where *B*_*absorbed*_ is the quantity (in mmol) of butyrate absorbed, *V*_*colon*_ and *S*_*colon*_ are the volume and surface area of the colon, respectively, [*B*] is the concentration of butyrate in the colon, and *γ*_*B*_ is the absorption rate of butyrate in mmol L^−1^h^−1^cm^−2^. We used two different estimates of *γ*_*B*_: the first corresponded to a published rate constant estimated using a dialysis bag technique (McNeil et al., 1978), and the second was obtained from our own measurements using a previously reported CaCo monolayer model system (Trapecar et al., 2019). We found that, assuming a transit time of 12 hours, these two rate constants have significantly different effects on the resulting dynamics: in the case of the dialysis bag constant (0.0019mmol L^−1^h^−1^cm^−2^), the majority of produced butyrate is excreted, while in the case of CaCo monolayer (0.091mmol L^−1^h^−1^cm^−2^), a significant quantity is absorbed (~40%) while the remainder is excreted (Fig. 4c). This indicates that, depending on the value of the absorption rate constant, the relationship between excreted stool SCFA quantities and quantities of SCFA produced and absorbed in the gut are not necessarily related.

Plotting the quantity of butyrate absorbed as a function of both absorption rate and transit time indicates that transit time is a significant variable controlling the quantities of SCFA absorbed by the gut in the orders of magnitude of the absorption rates considered (Fig. 4d). This relationship holds across different participants with different production rates of butyrate from the same input, related by a scaling factor. However, our model does not account for depletion of the fiber substrate, an event which measurements of stool concentrations of inulin (Fig. 4b and Fig. S3) suggest is likely to occur by the time the stool is excreted. Of course, slowing colonic transit time is therefore only an effective way of increasing absorbed SCFAs for as long as there remain SCFAs to absorb and fiber substrate to ferment. Nonetheless, it is clear from our model that, despite assuming equal absorption rate constants across participants and a transit time of 12h, different individuals absorb different quantities of an SCFA from a given fiber (Fig. 4e), mirroring their differences in production and overall MMP. Absorption is linearly related to the produced concentration, and changing absorption rate constant parameters only affects the scale of absorbed quantities but the relative ratios between individuals are unchanged.

## Discussion

In this study, we have used an *ex vivo* setup to measure the inter-individual differences in gut microbial SCFA production in response to different dietary fiber inputs. We have shown that there are significant differences in the MMPs of different individuals, i.e. differences in the capacity of their microbiota to ferment a given fiber substrate into a given SCFA. Moreover, we showed that MMP could to a certain extent be predicted from stool microbiota community composition, to a greater extent than can raw stool SCFA content. These data are consistent with a recent study where participants with particular microbiota compositions were more likely to yield higher butyrate concentrations in the stool after consuming resistant starch (Baxter et al., 2019).

In addition, we showed that the dominant features of an individual’s MMP are relatively stable through time, consistent with the fact that an individual’s MMP is related to their microbiota community composition. Though we did not study such cases, it is likely that treatment with broad-spectrum antibiotics or other extreme perturbations to the microbiota would have significant impact to an individual’s MMP, which concomitant implications to their physiology.

In order to explore the relationship between our *ex vivo* results and the implications for *in vivo* SCFA production and absorption, we sought to develop a phenomenological model of the process. We found that quantities of absorbed SCFAs mirrored the quantities produced in the different parameter regimes considered, though the absolute amounts varied significantly as a function of the epithelial absorption rate constants used in our model. This highlights the critical importance of this variable in understanding the relationship between stool SCFA quantities and *in vivo* production and absorption for clinical applications. It is therefore of great importance to the field to obtain accurate measurements of these rate constants in addition to quantitative descriptions of their behavior and the underlying kinetics of absorption at different concentrations of SCFAs and in different regions of the gut.

Our model also showed that increased colonic transit time results in significantly greater quantities of absorbed SCFAs while these remain at appreciable concentrations in the gut, or that there remains substrate to ferment. In our *ex vivo* experiments, as well as the *in vivo* inulin supplementation experiments used in Fig. 4b, the microbiota was given a highly accessible, powdered form of inulin as a substrate. Despite the fact that the majority of inulin is fermented by the microbiota both *ex vivo* and *in vivo* within 24h or by the time of passage, respectively, our *ex vivo* measurements indicate that this process occurs on the order of 12-24h in most participants at the considered input concentration (10g/L) (Fig. S3). Thus, it is likely that a less accessible form of fiber input (e.g. in the form of raw chicory root in the case of inulin) would take longer to ferment, in which case colonic transit time is an important variable in determining the quantities of SCFAs produced. Moreover, dietary fiber supplementation therapeutic strategies aiming to improve quantities of SCFAs produced would benefit from highly accessible forms of these fibers (purified powder), rather than grains, fruits and vegetables high in these quantities, which will likely take longer for the microbiota to ferment and therefore may result in a larger quantity of excreted, unfermented substrate.

Our model has several other limitations, which should be taken into account in the interpretation of our data. First of all, we do not consider fiber input concentration as a variable in the time evolution of our system, since we only considered a single concentration across participants (corresponding to significantly larger doses than those found in an ordinary diet) and assumed a linear production rate through time based on our *ex vivo* measurements (Fig. 1b). In addition, we assumed that epithelial absorption rates are identical across all participants. Moreover, it is unclear whether experimental production rates measured *ex vivo* are related to *in vivo* production in the gut.

Despite these limitations, our model suggests a high degree of robustness to the association between differences in MMPs and quantities of absorbed SCFAs across different individuals, despite the fact that varying the absorption rate constant had significant impact on both absorbed and excreted SCFA quantities. Importantly, depending on the actual magnitude of the absorption rate constant, the relationship between stool SCFA concentration and *in vivo* production and absorption can be completely non-informative. Stool SCFA concentration is related to the differential between production and absorption, the colonic transit time, and whether or not the entire fermentable substrate was consumed. For example, in a regime where production rate significantly exceeds absorption rate (as in both sets of absorption rate constant parameters considered in this study), stool SCFA is a function of the differential between the transit time (τ) and the time to depletion of the fermentable substrate (*t*_*1*_). If τ > *t*_*1*_, the stool SCFA concentration will mostly be a function of the time during which the stool was transiting but no further SCFAs were being produced (and luminal SCFAs were merely being absorbed). This is an important consideration that suggests that stool SCFA quantities may not be the relevant quantity of interest in the absence of knowledge of these other variables. Put differently, a lack of association between stool SCFA concentration and a clinical outcome variable is not necessarily indicative of a lack of involvement of microbial SCFAs in the disease process.

Taken together, our data indicate that a quantitative understanding of a patient’s MMP can inform personalized dietary supplementation strategies that aim to increase the production and resulting absorption of specific SCFAs in a patient’s gut. More broadly, they suggest a framework for modulating SCFA production in a patient through two separate but complimentary means: modification of dietary inputs as a function of their existing MMP, and modification of the underlying community composition of their microbiota towards a given MMP of interest.

## Acknowledgements

We are very grateful for the assistance provided by Prof. Linda Griffith and Jason Velazquez with the colonocyte monolayer experiments. In addition, we wish to thank Prof. Michael Fischbach for early insights and discussions regarding the ex vivo experimental setup. We also wish to thank Mary Delaney from the Harvard Digestive Disease Core for conducting the SCFA measurements, and Vicki Mountain for logistical support. This work was supported by a grant from the Rasmussen Family Foundation.

## Author contributions

This study was conceived and designed by T.G., L.T.T.N. and E.J.A. Raw data were collected by T.G., L.T.T.N. and X.Y. Final analyses were performed by T.G. and L.T.T.N. The paper was written by T.G., L.T.T.N. and E.J.A.

## Methods

### Human participants

Healthy human participants (14 females and 19 males) with ages ranging from 23 to 38 were consented to participate in this study under COUHES protocol number 1510271631.

### Stool sample processing and *ex vivo* setup

Fresh stool samples were collected and weighed before being transferred to anaerobic conditions, in which they were diluted in reduced PBS containing 0.1% L-Cysteine to a ratio of 1g/5ml. Samples were homogenized into a slurry before being aliquoted into 96-well plates. Samples of the unaltered slurry were taken for a baseline sample, after which inulin, pectin, and cellulose were added from stock solutions to final slurry concentrations of 10g/L, 5g/L and 20g/L, respectively. Concentrations of inulin and pectin were determined based on the maximum stock concentration we were able to obtain in which the dietary fiber was fully dissolved. Four conditions were measured: inulin, pectin, cellulose and control (no spike-in). The samples were incubated anaerobically at 37°C, and two biological replicates (two different wells in the 96-well plates) were collected at each time point. Samples were collected after 2 and 4 hours from all participants and sent out for SCFA quantification on a GC-FID. Linear production rate was measured between 2h and 4h because this allowed us maximum accuracy in measuring the time interval without introducing artefacts due to delays between conditions introduced during setup. Results obtained from slurry in large flasks on a shaker were in good agreement with data obtained in a 96-well plate format.

### Short-chain Fatty acid measurements

Chromatographic analysis was carried out using an Agilent 7890B system with a flame ionization detector (FID) (Agilent Technologies, Santa Clara, CA). A high resolution gas chromatography capillary column 30m x 0.25 mm coated with 0.25 μm film thickness was used (DB-FFAP) for the volatile acids (Agilent Technologies) and a high resolution gas chromatography capillary column 30m x 0.25 mm coated with 0.50 μm film thickness was used (DB-FFAP) for the nonvolatile acids. Nitrogen was used as the carrier gas. The oven temperature was 145°C and the FID and injection port was set to 225°C. The injected sample volume was 1μL and the runtime for each analysis was 12 minutes. Chromatograms and data integration was carried out using the OpenLab ChemStation software (Agilent Technologies).

For chromatographic analyses, a volatile acid mix containing 10mM of acetic, propionic, isobutyric, butyric, isovaleric, valeric, isocaproic, caproic, and heptanoic acids was used (Supelco CRM46975, Bellefonte, PA). A standard stock solution containing 1% 2-methyl pentanoic acid (Sigma-Aldrich St. Louis, MO) was prepared as an internal standard control for the volatile acid extractions.

Samples were kept frozen at −80°C until analysis. The samples were removed from the freezer and allowed to thaw. A sample of the raw fecal material was transferred to a 2mL tube, the weight of the fecal material was determined and 1.5 mL of HPLC water was added to each sample. The samples were vortexed for 5 minutes until the material was homogenized. The pH of raw fecal suspension and the thawed fecal slurry samples was adjusted to 2-3 with 50% sulfuric acid. The acidified samples were kept at room temperature and vortexed for 10 minutes. The samples were centrifuged for 10 minutes at 5000g. 1000uL of the clear supernatant was transferred into a glass tube with a PTFE faced rubber lined screw cap for further processing. 50uL of the internal standard (1% 2-methyl pentanoic acid solution) and 1ml of ethyl ether anhydrous were added to the volatile samples. The tubes were mixed end over end for 10 minutes and then centrifuged at 2000g for 1 minute. The upper ether layer was transferred to an Agilent sampling vial for analysis.

For quantification of SCFAs, 1ml of each of the standard mixes were used and processed as described for the samples. The retention times and peak heights of the acids in the standard mix were used as references for the sample unknowns. These acids were identified by their specific retention times and the concentrations determined and expressed as mM concentrations per gram of sample for the raw fecal material and as mM concentrations per mL of fecal slurry.

### 16S rRNA sequencing

For DNA extraction, the MoBioPowersoil 96 kit (now Qiagen Cat No./Id: 12955-4) was used with minor modifications. All samples were thawed on ice and 250uL of the 5x dilution fecal slurry from the *ex vivo* assay from each sample were transferred to the Mobio High Throughput PowerSoil bead plate (12955-4 BP) for sample loading steps. We then proceeded through the extraction procotol on the same day following the manufacturer’s instructions.

Paired-end Illumina sequencing libraries were constructed using a two-step PCR approach targeting 16S rRNA genes V4 region, previously described by Preheim et al. (Preheim et al., 2013). All paired-end libraries were multiplexed into lanes (at maximum 200 individual samples pooled per lane) and sequenced with paired end 150 bases on each end on an Illumina MiSeq platform.

### Inulin-specific ELISA

We use BioPAL’s inulin immunoassay kit (BioPAL Worcester MA) to measure inulin concentration in filtrate from stool samples and followed the manufacturer’s protocol. For stool samples that came from the *ex vivo* assay, they were already diluted 5x and needed to be centrifuged (10’000xg for 2 minutes) before passing the supernatant through a 0.2 μm syringe filter (Pall Corporation, Port Washington NY). For stool samples that came from the *in vivo* diet study (Gurry et al., 2018), they were thawed on ice and homogenized to a 5x dilution fecal slurry in PBS buffer before being spun down and filtered similarly to the *ex vivo* samples. The fecal filtrates were stored at −80°C, always thawed on ice, and diluted to an appropriate dilution to be in the detection range of the ELISA Inulin kit. For reference, the *ex vivo* inulin conditions needed to be diluted at least 1’000-fold while the *ex vivo* control or *in vivo* samples only needed to be diluted at 10-fold.

### Gut monolayer SCFA absorption measurements

We used a gut monolayer system without the immune component, where the gut monolayer was prepared as described previously by WLK Chen et al. (Chen et al., 2017b; Trapecar et al., 2019). Briefly Caco-2 or C2BBe1 along with HT29-MTX-E21 cells (Sigma) were seeded onto rat-tail collagen I-(corning 354236) coated Transwell inserts (Corning 3460) in a 9:1 ratio (1 x 105 cells/cm2) in 500 uL seeding medium. The top and bottom compartments of the Transwell Plate were fed with 500uL and 1.5mL of seeding medium. The medium was changed every 2-3 days. After 20 days of gut insert maturation, the cell monolayer was ready to be used.

Barrier integrity was quantified by TEER (TransEpithelial Electrical Resistance) using the EVOM2 and the Enfohm-12 (World Precision Instruments) at 37°C. We measured cell monolayer integrity at the beginning and the end of the 24h long experiment. At the in-between time points, we surveilled the cell monolayer’s (dis)continuity under the microscope.

SCFAs were added to the mammalian cell media in their salt forms: Sodium Butyrate (Sigma, B5887), Sodium propionate (Alfa Aesar, A17440) and Sodium acetate (Sigma, S2889). They were added to the gut cell media on the apical side while the basal side was left unchanged, and the experiment was run for 24 hours. At each of the time points (0, 2, 4, 8 and 24h) we collected 100μL from the apical side and 200μL from the basal side from each gut cell Transwell insert (previous experiments done by WLK Chen et al. have shown that the resulting reduction in volume did not affect the cell culture). Collected media from apical and basal sides were stored at −80°C until analyzed for their SCFA contents. Each condition was run in triplicates except for the controls.

We ran 5 different concentrations of SCFA combinations. The Butyrate:Propionate:Acetate ratio (1:1:5, respectively) was kept constant across the different conditions. Concentrations tested were 40mM, 20mM, 10mM, 5mM and 2mM of Butyrate, with corresponding amounts of Propionate and Acetate according to the aforementioned ratio. The controls were either single SCFA (5mM Butyrate, 5mM Propionate or 25mM Acetate) or without added SCFA. At 24h, all the cells monolayers were washed, lysed and kept frozen at −80°C for further analysis if needed.

### 16S rRNA sequencing analysis

Raw paired-end 16 S rRNA Illumina sequencing reads were merged, demultiplexed, and quality trimmed with a cut-off of Q = 25 using usearch8, before being trimmed to a common length of 226 bases. Dereplicated reads were then clustered into OTUs to 97% identity using UPARSE (Edgar, 2013). OTU centroids were assigned a taxonomy using the RDP classifier using an uncertainty cut-off of 0.5 (Cole et al., 2005).

## Machine learning

Random Forest Classifiers (RFCs) were built using the scikit-learn Python package. 5-fold cross-validation was used to construct an average Receiver Operator Characteristic (ROC) from which to compute an AUC.

## Supplementary Information

### Model of microbial SCFA production

Figure S9a illustrates the steps in the degradation of dietary fibers into SCFAs that involve the microbiota. In this framework, distinct equations describing the time-evolution of the concentration of each chemical species exist for each type of monomer *M*, intermediate *P*, and SCFA. In reality, the system is sparse, as the rate constants *π*_*i1*_, *π*_*j2*_, and *π*_*k3*_ are zero or close to zero in the vast majority of OTUs due to the fact that only a small subset of OTUs have the necessary biochemical capabilities. For a given concentration of dietary fiber [*F*], the combination of all contributing equations to a given rate of production of a specific 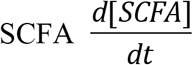 can be summarized by a single parameter, 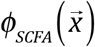, which is a function of the composition of the microbiota (Fig. S9b). Thus, in *ex vivo* conditions,

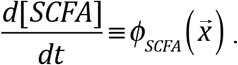

### Estimation of absorbed SCFAs across the colon’s epithelium

We approximate the colon as a cylinder 3cm in diameter. This corresponds to a circumference of approximately 9.42cm. Thus, each 1cm-thick cross-sectional segment of the colon has a surface area of approximately 9.42cm^2^, which we round up to 10cm^2^. Estimating the length of the colon at 150cm, we reach a total surface area of approximately 1,500cm^2^. Assuming the colon is filled with stool, we reach a total internal stool volume of 1,060cm^3^, which we approximate as 1L.

The absorption rate constant of a given SCFA estimated using the CaCo cell monolayer experiments, *ϒ*_*SCFA*_, is in units of mol cm^−2^ h^−1^L^−1^. Thus, the absorption rate of the entire colon is 1,500*ϒ_SCFA_* mol h^−1^L^−1^ of each SCFA. Based on our approximation of 1L of total stool, this amounts to an absorption rate of approximately 1,500*ϒ*_*SCFA*_ mol h^−1^ for each SCFA.

### Comparison between *ex vivo* and *in vivo*

We next sought to determine whether changes in SCFAs or community structure observed under *ex vivo* conditions were in agreement with what could theoretically be measured *in vivo*. For this purpose, we turned to a dataset from a previous study in which participants were placed on a fixed diet consisting entirely of a fiber-impoverished, liquid, nutritional meal supplement for a period of six days (Gurry et al., 2018). In the latter three days, participants were randomized to a spike-in, to be consumed at a prescribed dose daily against the constant liquid diet background. These spike-ins included inulin, pectin, and cellulose, and used the exact same sources of these three fibers, providing us with an ideal comparison dataset.

Under *ex vivo* conditions, we observed complete degradation of the inulin bolus in certain participants (Fig. 3a). We sought to determine whether a similar extent of degradation could be observed *in vivo* in participants consuming an inulin bolus at similar concentrations. Residual inulin was therefore quantified from the stool using an inulin-specific ELISA assay. We find that, as expected, no detectable inulin can be found in the stool of participants on days 1, 2 and 3, but that inulin is detectable on days after which inulin was consumed in certain stool samples (Fig. 4b). However, the inulin detected on these days accounts for a tiny minority of the total inulin consumed (10g/day), which assuming a total daily stool volume of 1L, equates to approximately 10g/L, the concentration used in the *ex vivo* experiments. This suggests that the majority of accessible inulin is also consumed *in vivo* in the typical stool passage time following ingestion. Moreover, since 10g is a significantly larger dose of inulin than would ordinarily be consumed in a typical diet, we can conclude that the majority of inulin consumed in an ordinary diet in a form similarly accessible to the inulin powder used for these experiments is fermented *in vivo*. Of course, it is likely that inulin from natural sources and typical dietary fiber sources do not contain the fiber in as accessible form as the purified form used in these experiments.

In order to assess the extent to which the *ex vivo* conditions result in the growth of different organisms compared to *in vivo*, we recruited the same participants from the *in vivo* study that had consumed inulin as a spike-in, performed an *ex vivo* experiment, and directly compared the 16S rRNA data between the two so the comparison could be made within the same person. More specifically, we compared OTUs changed *in vivo* from day 3 to day 6, or changed *ex vivo* from *t*=0 to *t*=24h, and plotted their trajectories next to each other (Fig. S10). Comparing two separate OTUs of the same butyrate-producing genus, *Blautia*, we find that participant OTU denovo29 blooms in participants G and V *in vivo* but in participant F *ex vivo*, while OTU denovo78 blooms in all participants *ex vivo* but does not change appreciably *in vivo*. Thus, these data argue that *ex vivo* conditions may introduce artefacts in growth rates of different bacteria compared to the gut, further arguing for measurement in the first hours of the experiment while the stool community remains relatively unchanged.

**Figure S1.**
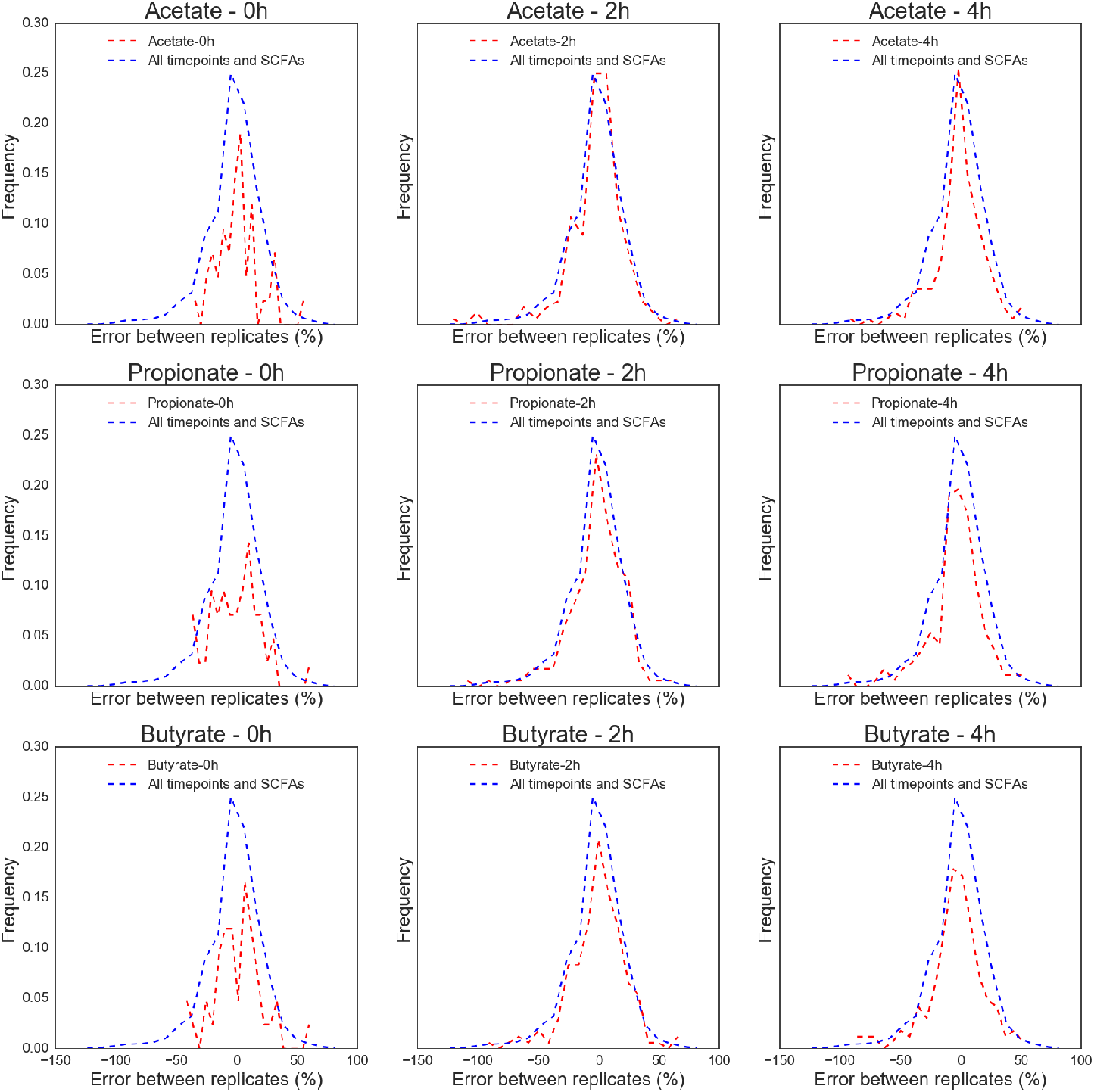
Percentage difference in SCFA quantification by GC-FID between biological replicates, broken down by timepoint and SCFA. The overall distribution, across all participants, timepoints, and SCFAs, is plotted in blue, while the individual distributions for each timepoint and SCFA are shown in red.

**Figure S2.**
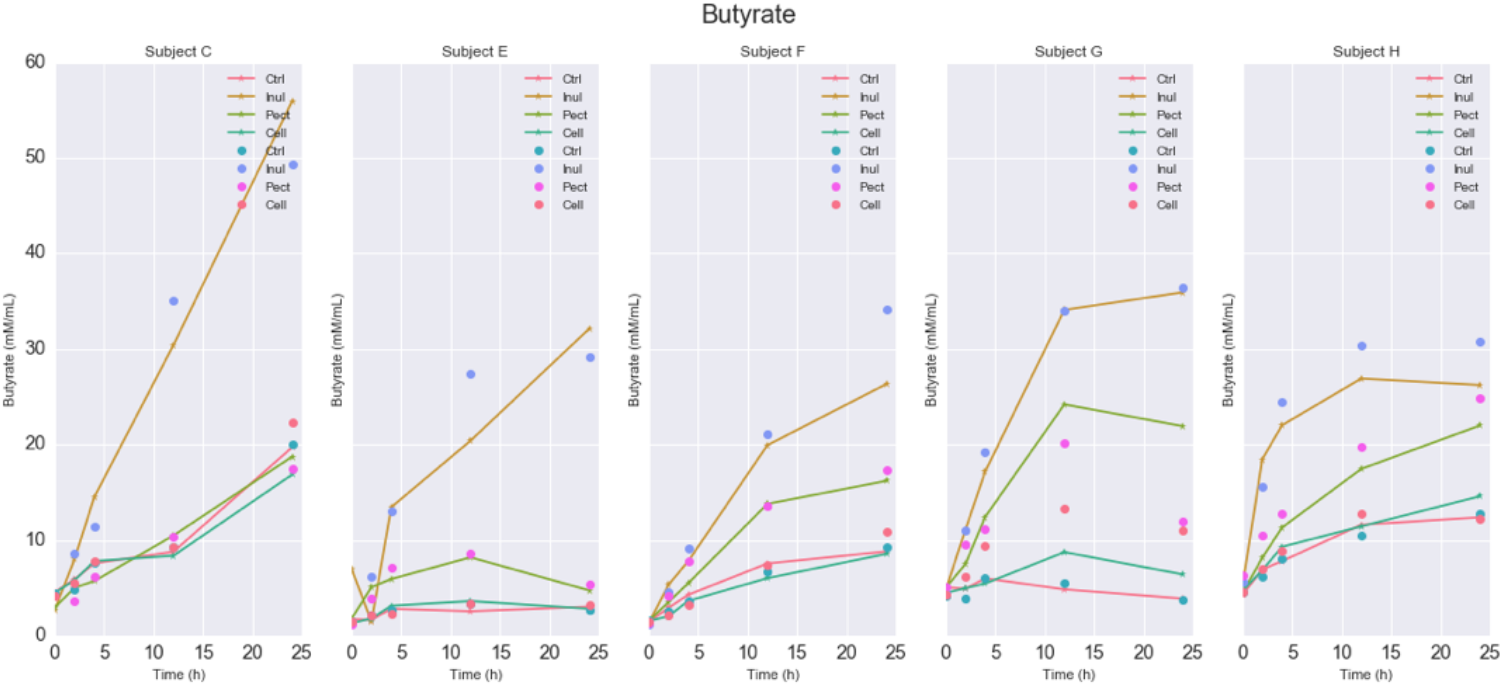
Individual 24h trajectories for butyrate in five pilot participants. The different conditions are shown in different colors.

**Figure S3.**
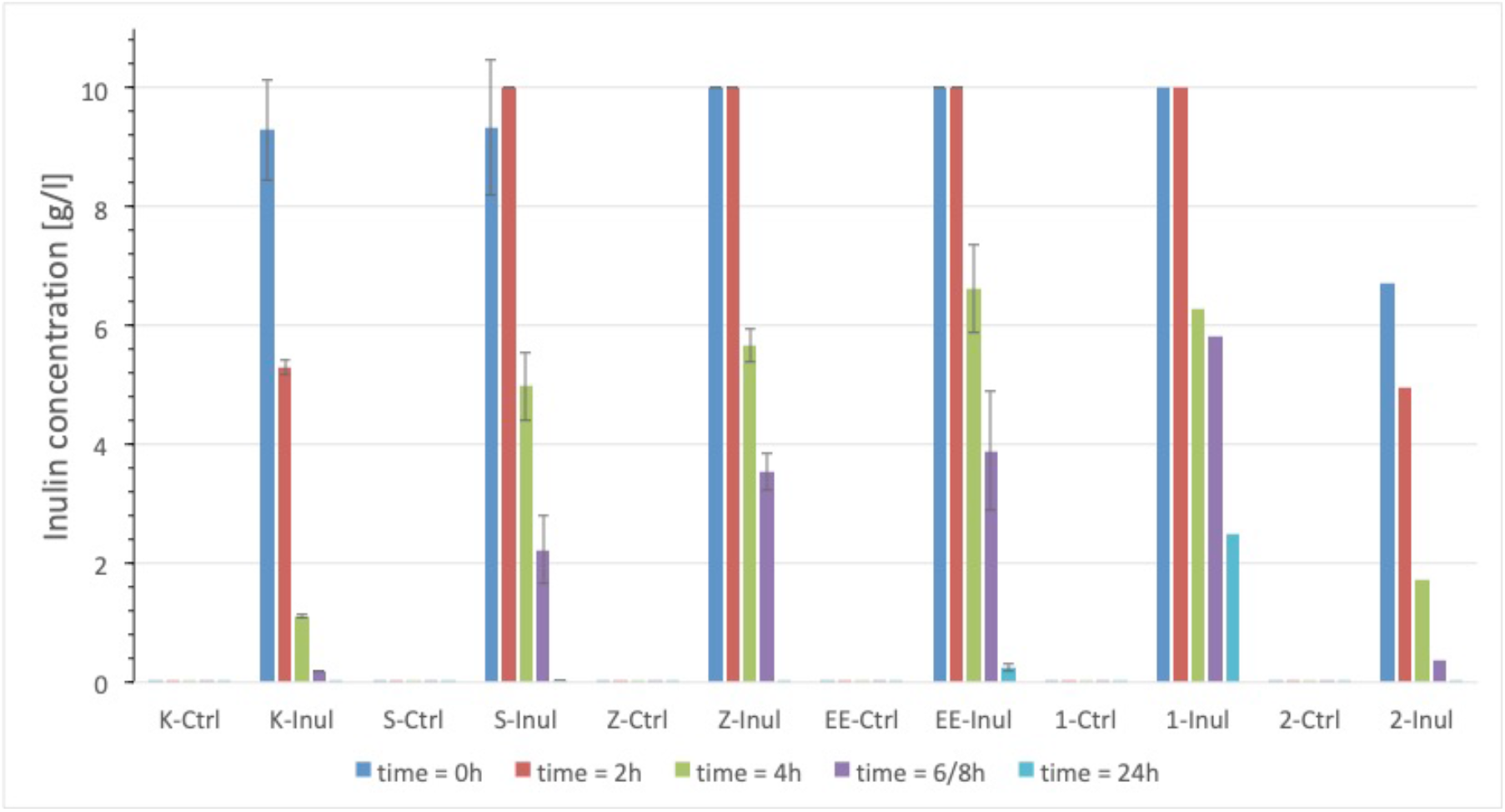
Concentrations of inulin measured at *t* = 0, 2h, 4h, 6h and 24h in six separate donors (donor IDs K, S, Z, EE, 1 and 2), determined using an inulin-specific ELISA assay.

**Figure S4.**
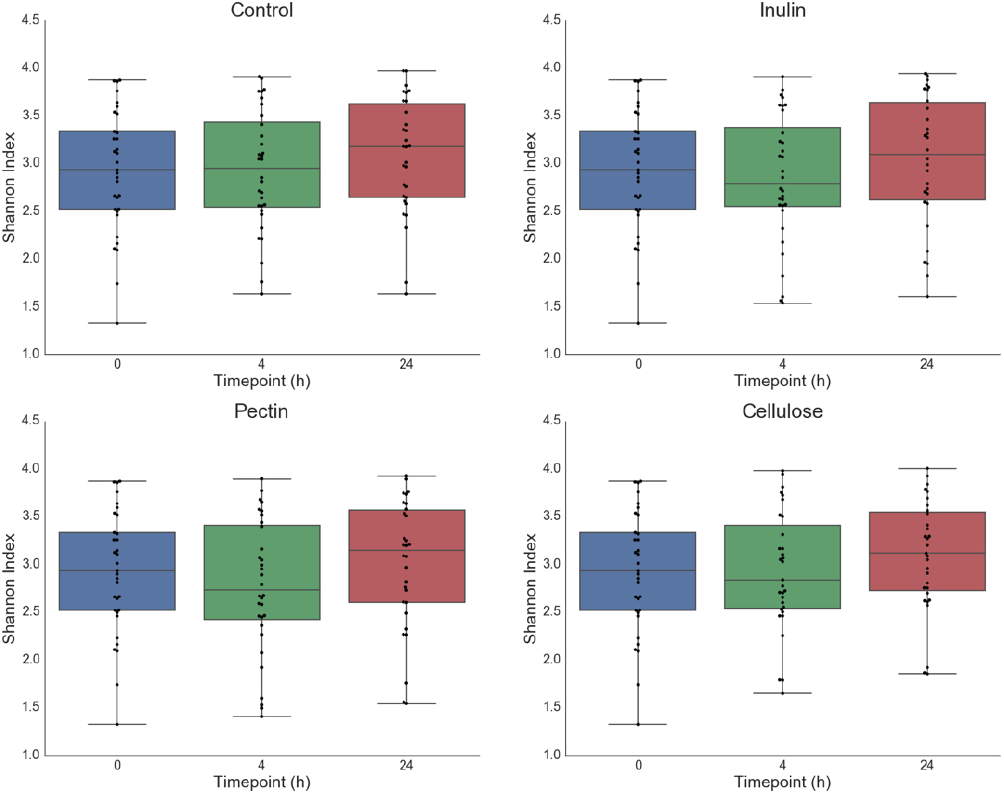
Shannon diversity index of 16S rRNA communities in each condition and each timepoint (0h, 4h, 24h).

**Figure S5.**
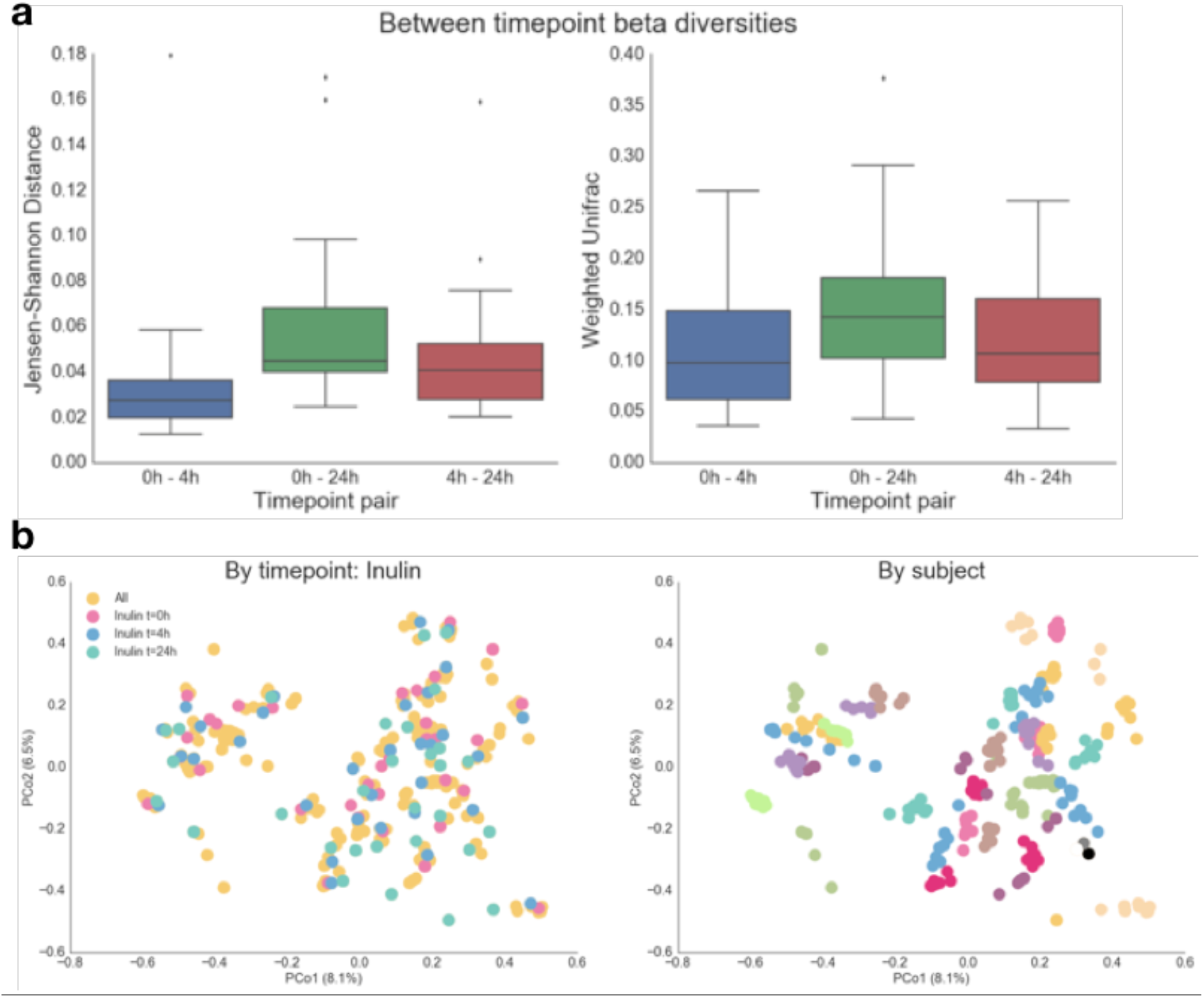
(a) Jensen-Shannon Distances (left) and Weighted Unifrac (right) beta-diversities between timepoints at the level of 16S rRNA (0-4h, 0-24h, 4-24h). (b) Multidimensional scaling analysis of all 16S rRNA samples, colored by timepoint (left) and by participant (right).

**Figure S6.**
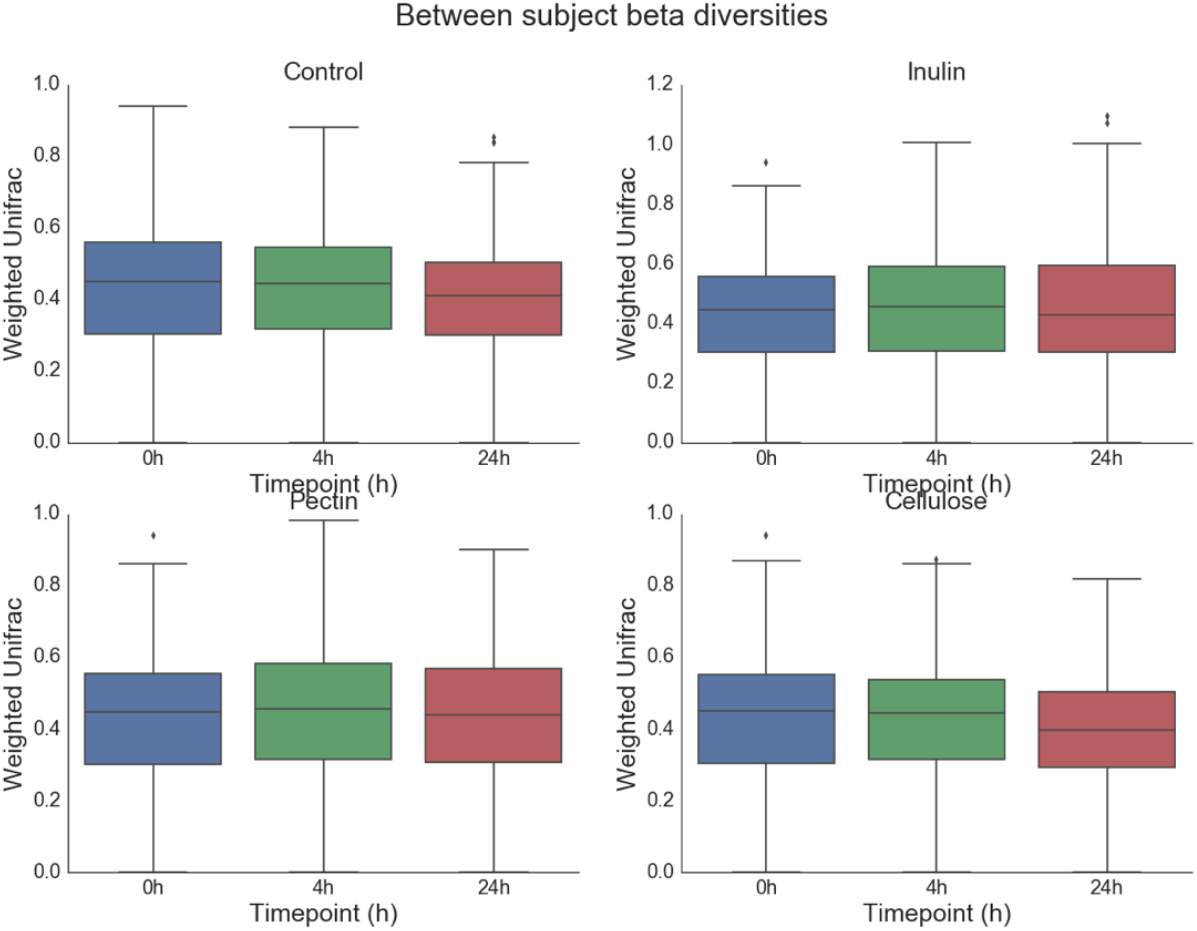
Between participant Weighted Unifracs at 0h, 4h and 24h.

**Figure S7.**
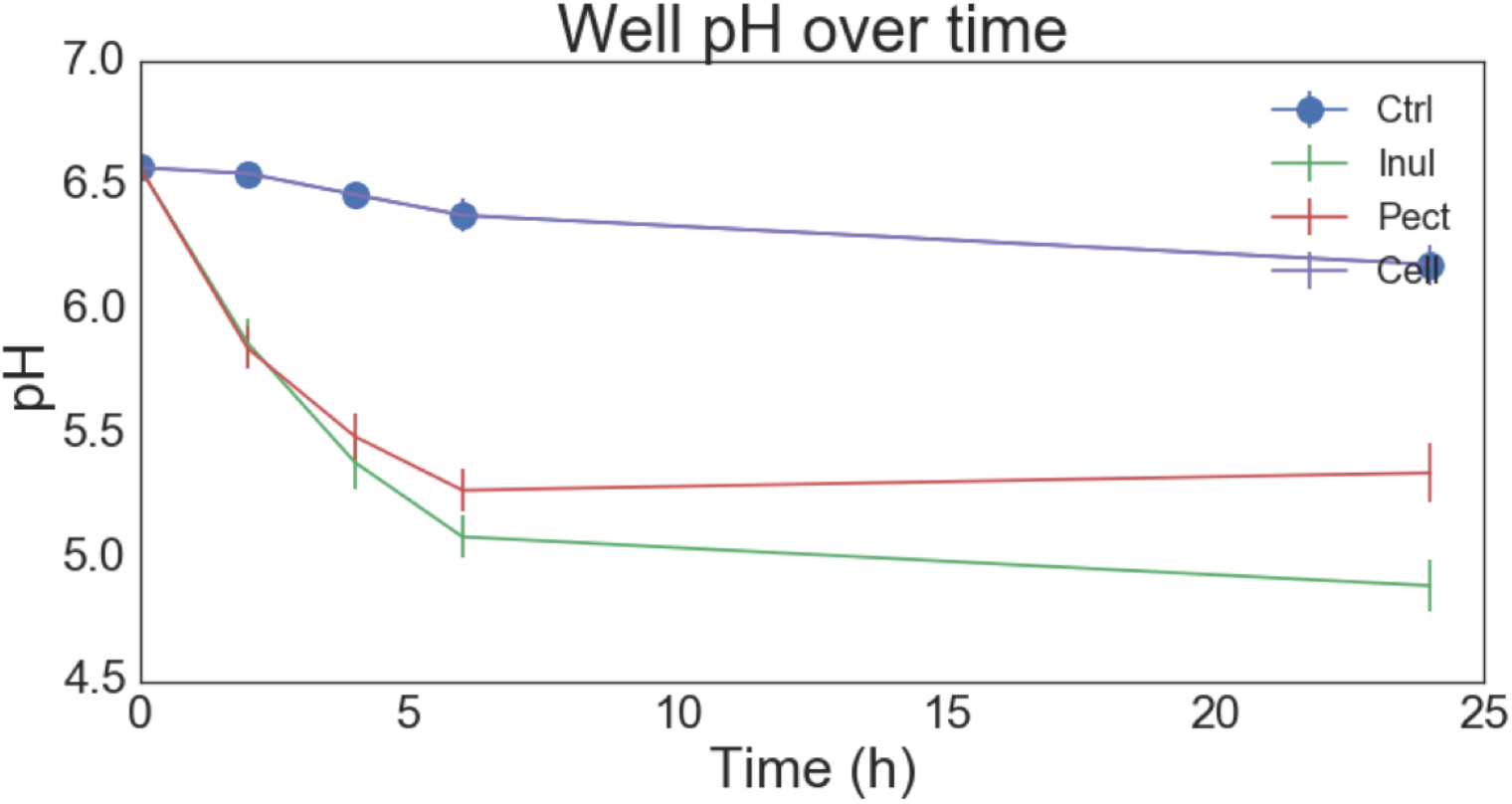
pH of the slurry over time, measured across all participants in the study.

**Figure S8.**
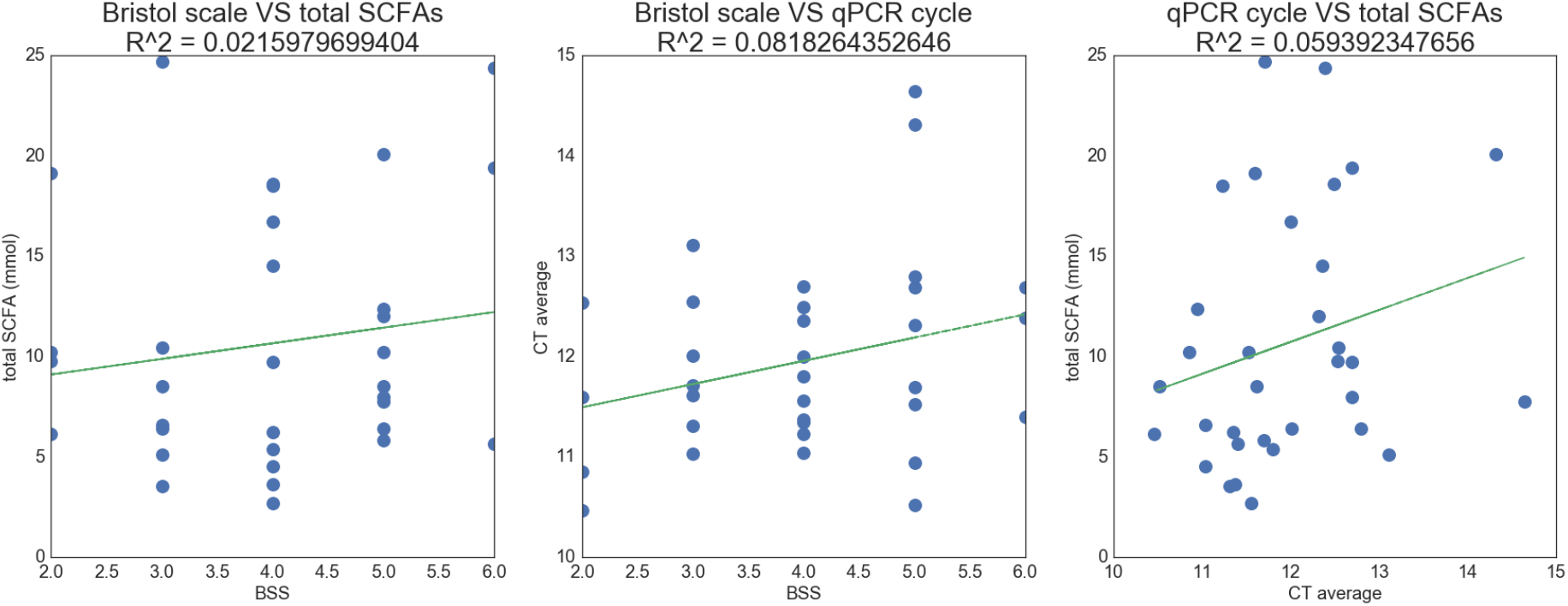
Pairwise linear regressions between between Bristol score of the sample, total SCFAs produced, and qPCR amplification cycle (C_T_ values).

**Figure S9.**
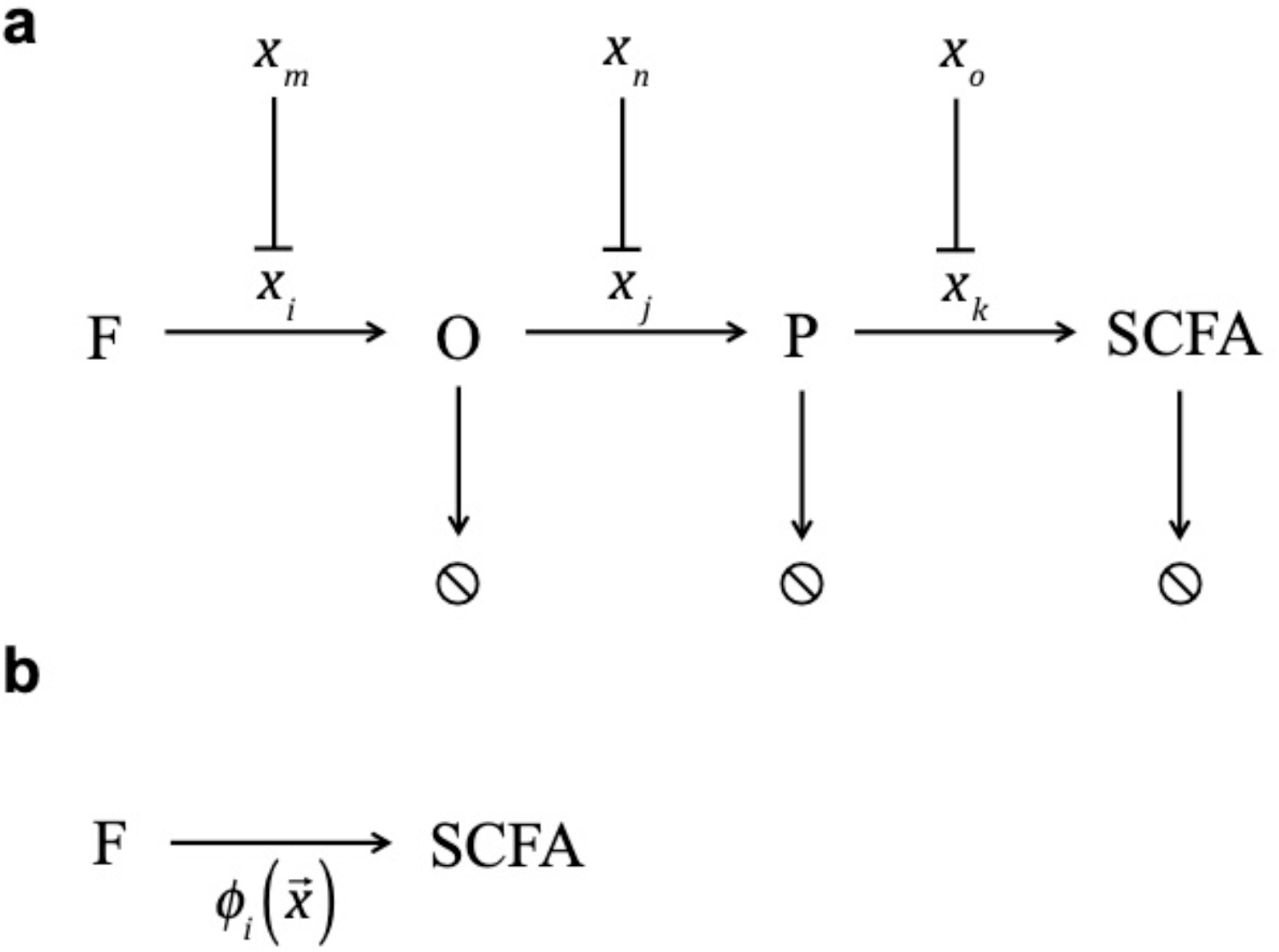
Microbial SCFA production proceeds through the cooperation of different members of the microbiota. (a) Schematic illustrating the generic steps involved in dietary fiber degradation. A bacterial OTU *i* of relative abundance *x*_*i*_ hydrolyses the polysaccharide (dietary fiber) *F* into oligosaccharides *O*. These are then fermented into a reduced intermediate *P* by OTU *j* with relative abundance *x*_*j*_. Finally, *P* may be further fermented to an SCFA by OTU *k* with relative abundance *x*_*k*_. In addition, the ability of an OTU to carry out a given reaction can itself be inhibited by a separate OTU (e.g. *x*_*i*_ is inhibited by *x*_*o*_). (b) Bulk measurement of the overall production rate of a given SCFA, *ϕ*_*SCFA*_(**x**), which is itself a function of the composition of the stool microbiota, ***x***, and corresponds to the quantities measured using our *ex vivo* experiments.

**Figure S10.**
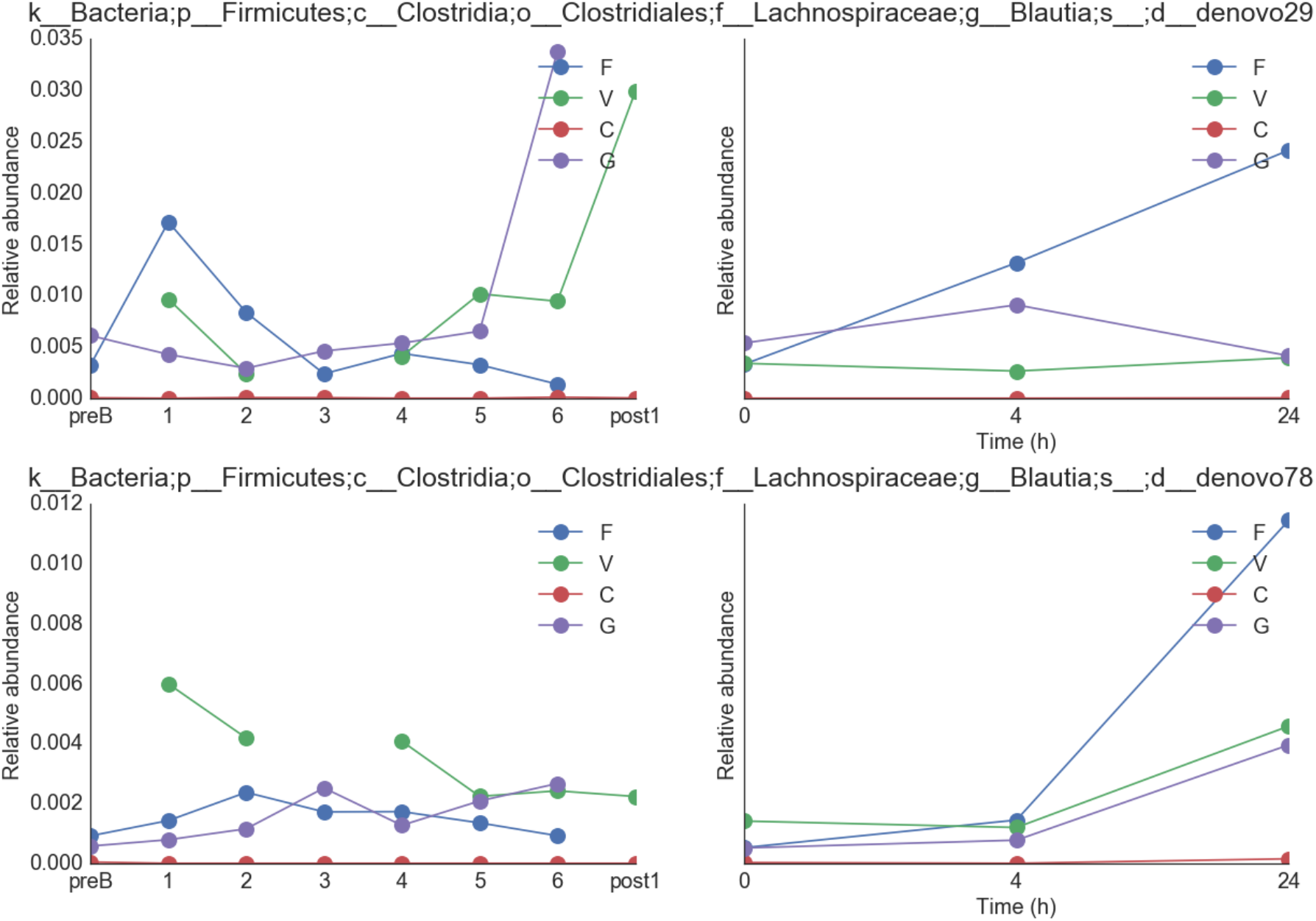
Comparison in growth dynamics of two different OTUs of the genus *Blautia* in the same participants *in vivo* and *ex vivo*.

